# Tokenizing single-cell transcriptomes as a native language for large language models

**DOI:** 10.1101/2025.10.22.684047

**Authors:** Chuxi Xiao, Yuang Ding, Haiyang Bian, Yixin Chen, Lei Wei, Xuegong Zhang

**Affiliations:** MOE Key Laboratory of Bioinformatics and Bioinformatics Division, BNRIST, Department of Automation, Tsinghua University, Beijing, China; Tongyi Lab, Alibaba Group, Beijing, China; Institute of Science and Technology for Brain-inspired Intelligence, Fudan University, Shanghai, China; School of Life Sciences and School of Medicine, Center for Synthetic and Systems Biology, Tsinghua University, Beijing, China

## Abstract

Large language models (LLMs) can process diverse forms of information once they are represented as tokens in a shared sequence space. However, single-cell transcriptomes remain a foreign modality to LLMs because they are continuous, high-dimensional molecular profiles rather than discrete linguistic units. Here, we propose CellTok, a tokenized single-cell language modeling approach that converts transcriptomic profiles into compact cellular token sequences and incorporates them into the vocabulary of a pretrained LLM. By representing cells as native tokens, CellTok enables cellular measurements, textual instructions, biological context, and multi-cell populations to be jointly processed within the same autoregressive modeling framework. Across diverse tasks, CellTok enable LLMs to recognize individual cells, interpret homogeneous and heterogeneous cell populations, infer disease-associated cellular states, predict cell–cell communication, model developmental trajectories, and generate cellular states. Moreover, prompt-based experiments show that providing appropriate biological context improves performance, indicating that CellTok can leverage LLM knowledge and contextual reasoning to support cellular data interpretation. These results demonstrate that single-cell transcriptomes can be transformed from a foreign molecular modality into a native language for LLMs, establishing a unified interface for modeling cells, populations, and biological knowledge in a shared token space.

## Introduction

Large language models (LLMs) have shown remarkable capacity to process, compose, and reason over information expressed as tokens^1,2^. Their success across natural language, code, structured records, and multimodal inputs suggests that token-based sequence modeling can provide a general interface for representing diverse forms of knowledge^3–5^. However, this interface has not yet extended to single-cell biology. Single-cell transcriptomes are continuous, high-dimensional molecular measurements rather than discrete linguistic units. To an LLM, a cell is therefore still largely a foreign language: it can be described by text, summarized by a label, or projected into an embedding space, but its molecular state cannot be directly read, composed, predicted, or generated as part of the model’s native token language.

Recent large cellular models (LCMs) have substantially advanced the representation of cellular states by learning from large-scale transcriptomic data^6–13^. Many of these models are inspired by language modeling, but they are not native languages for general-purpose LLMs. As a result, cellular states remain external to the token space of LLMs and cannot be composed with text or conditioned on by prompts. Recent efforts have therefore explored more direct uses of LLMs in single-cell analysis^14–19^, for example by serializing transcriptomic profiles as gene-level textual sequences^14^, or aligning continuous cellular embeddings with textual representations through cross-modal learning^15^. These studies suggest that LLM knowledge can help interpret cellular data, but the connection between cells and LLMs remains indirect. In these formulations, cellular information is first converted into textual summaries, projected features, or task-specific representations, rather than being provided to the LLM as native cellular token sequences.

This indirect interface becomes limiting when cellular analysis requires contextual reasoning over cells. Many central problems in single-cell biology do not concern isolated transcriptomes alone, but how cells are grouped into populations, how cells communicate with one another, and how cellular states are ordered along developmental or disease processes^20–22^. These problems require cells to be represented as compositional units that can be arranged with other cells, interpreted under task-specific instructions, and connected to biological knowledge. This is precisely the type of contextual reasoning for which LLMs are well suited, but it requires cells to be part of the model’s token space rather than external vectors or textual summaries. We therefore reasoned that cellular tokenization could provide a native interface between single-cell transcriptomes and LLMs. If transcriptomic profiles can be converted into compact discrete cellular tokens and incorporated into the vocabulary of a pretrained LLM, then cells can be processed in the same sequence space as language, allowing cellular measurements, multi-cell contexts, textual prompts, and biological knowledge to be modeled within a unified autoregressive framework.

Here, we propose CellTok, a tokenized single-cell language modeling approach that transforms transcriptomic profiles into native cellular tokens for LLMs. CellTok first learns a compact vocabulary of cellular states using a vector-quantized autoencoding strategy, representing each cell as a short sequence of discrete tokens. These cellular tokens are then added to the vocabulary of a pretrained LLM and modeled together with text tokens under an autoregressive objective. This design allows individual cells, cell populations, task instructions, and biological context to be represented within a shared token space.

We evaluate CellTok across tasks that test whether LLMs can read and reason over cellular data. CellTok enables LLMs to recognize cellular identities, interpret homogeneous and heterogeneous cell populations, infer disease-associated cellular states, predict cell–cell communication, model developmental trajectories, and generate cellular states under specified contexts. We further show that biologically informative prompts improve performance, suggesting that CellTok can use LLM knowledge and contextual reasoning to support cellular interpretation. Together, these results demonstrate that single-cell transcriptomes can be transformed from a foreign molecular modality into a native language for LLMs, establishing a route toward cellular analysis in which cells, populations, task instructions, and biological knowledge are modeled in a shared sequence space.

## Results

### CellTok learns compact cellular tokens for native sequence modeling within LLMs

We developed CellTok, an approach that transforms single-cell transcriptomes into discrete token sequences and enables their native modeling within large language models (LLMs) through an early-fusion design (**Fig. 1A**). At the core of CellTok is a learned tokenization mechanism that represents each cell’s gene expression profile as a compact sequence of discrete tokens. Inspired by vector-quantized variational autoencoders (VQ-VAE)^23^, we employed an encoder–quantizer–decoder architecture to compress high-dimensional transcriptomic profiles into a short sequence of discrete codes designed to preserve biologically relevant expression patterns. These learned codebook vectors are incorporated directly into the vocabulary of a pretrained LLM (Qwen 2.5 7B^24^) as cell tokens, where they participate in autoregressive modeling in the same manner as conventional text tokens. Unlike late-fusion approaches, this early-fusion design allows cellular tokens to serve directly as conditioning inputs, contextual elements, and prediction targets, and enables interactions between cellular and linguistic information throughout the model.

**Fig. 1.**
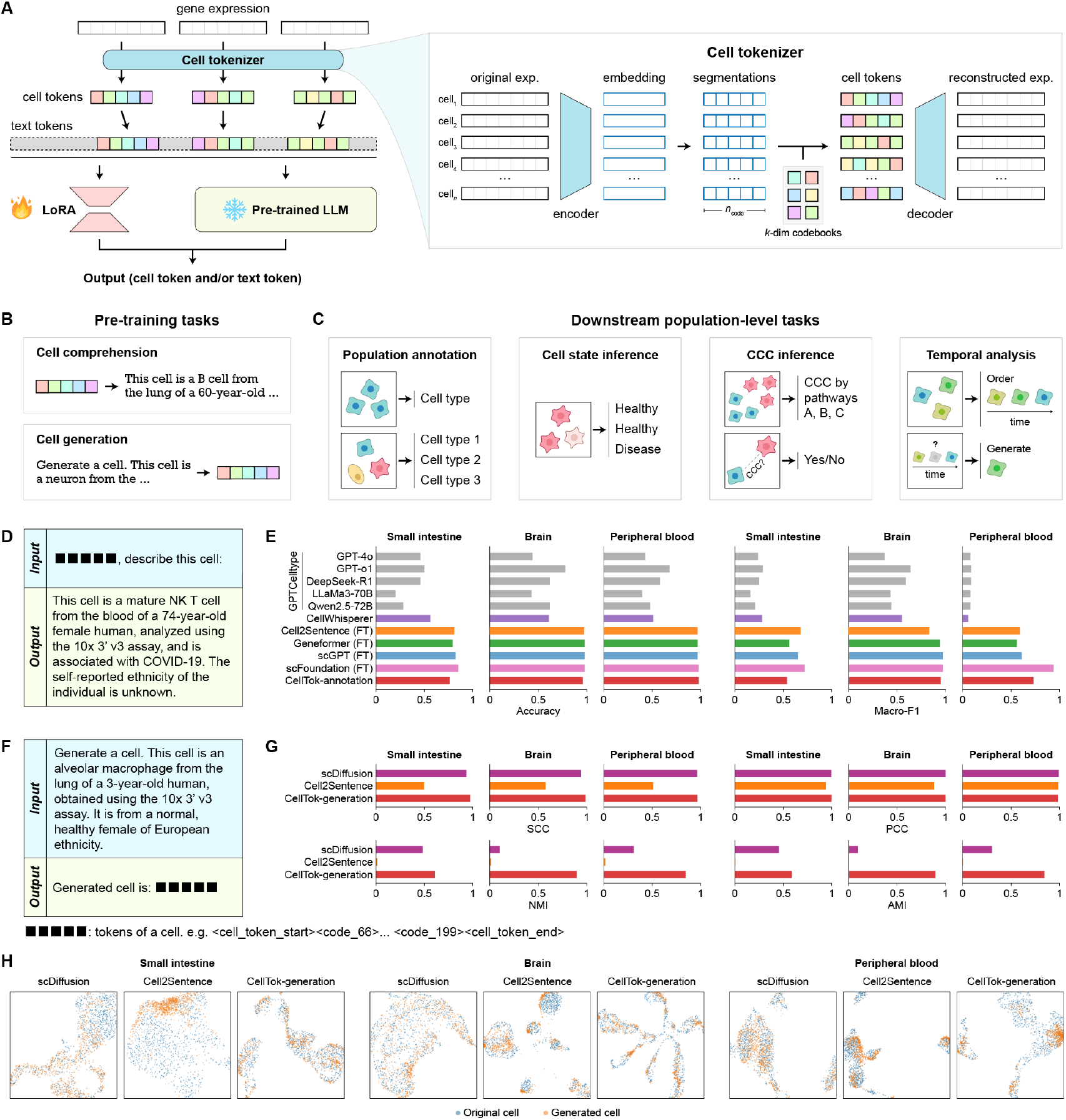
Overview of CellTok. (A) Architecture of CellTok. Single-cell gene expression profiles are compressed into discrete cell tokens using an encoder–quantizer–decoder mechanism. These learned tokens are directly integrated into the vocabulary of a pretrained large language model (LLM) via an early-fusion design, enabling joint autoregressive modeling of cellular and textual information. (B) Dual pretraining tasks of CellTok. CellTok is trained using a cell comprehension task (generating textual descriptions conditioned on cell tokens) and a cell generation task (predicting cell tokens conditioned on textual descriptions). (C) Downstream population-level tasks of CellTok. (D) Schematic of the cell type annotation task of an individual cell. (E) Performance of CellTok-annotation against baseline methods on individual-cell annotation. (F) Schematic of the conditional generation task. (G) Performance of CellTok-generation against baseline methods on conditional generation. (H) UMAP visualization of original and generated cells. SCC: Spearman correlation coefficient; PCC: Pearson correlation coefficient; NMI: normalized mutual information; AMI: adjusted mutual information.

To connect cellular tokens with biological language, we trained CellTok with two complementary pretraining objectives (**Fig. 1B**). A cell comprehension task conditions on cell tokens to generate textual descriptions of cellular states, linking discrete transcriptomic representations to biological concepts. Conversely, a cell generation task conditions on textual descriptions to generate corresponding cell tokens, which can be decoded into transcriptomic profiles by the decoder. These objectives unify understanding and generation across modalities within a single autoregressive framework. We trained CellTok-base on 68 million high-quality single-cell profiles curated from CELLxGENE^25^ and paired the profiles with textual descriptions generated using GPT-4o^26^ (**Fig. S1, Methods**). Consistent with a scaling behavior of language models, larger LLM backbones improved CellTok performance in our backbone comparison (**Fig. S1E**), indicating that cellular token modeling is compatible with model scaling. This training provides CellTok-base with a general cellular token interface that can subsequently be adapted to cell-level and population-level tasks (**Fig. 1C**).

After training, we first examined whether cellular tokens preserve enough biological information for individual-cell interpretation. We evaluated CellTok on cell type annotation of individual cells across multiple datasets held out from CellTok-base pretraining, including the small intestine^27^, brain^28^, and peripheral blood^29^ datasets (**Fig. 1D**). Given the tokenized representation of a cell, CellTok generated a textual description, from which the predicted cell type was extracted and compared with the ground-truth label. After task-specific fine-tuning of CellTok-base, CellTok (CellTok-annotation) achieved performance competitive with LCMs (Geneformer^9^, scGPT^10^, and scFoundation^11^) as well as LLM-based analysis methods (CellWhisperer^15^ and Cell2Sentence^14^) across multiple metrics, and notably outperformed substantially larger general-purpose LLMs (GPT-4o^26^, GPT-o1^30^, DeepSeek-R1^31^, LLaMA 3 70B^32^, and Qwen 2.5 72B^24^) prompted for scRNA-seq analysis following the GPTCelltype framework^17^ (**Fig. 1E, Methods**). These results indicate that CellTok retains cell-identity information in a discrete token format that can be interpreted through the autoregressive language modeling interface.

We next asked whether the same token interface supports generation of cellular states. Conditioning on textual descriptions, CellTok can generate sequences of cell tokens that are decoded into gene expression profiles (**Fig. 1F**). A fine-tuned generative variant (CellTok-generation, fine-tuned with the same held-out datasets) produced synthetic cells that closely match real transcriptomic profiles, substantially outperforming Cell2Sentence^14^ and achieving performance comparable to scDiffusion^33^ across multiple metrics (**Fig. 1G–H, Methods**). This demonstrates that the learned cellular tokens capture enough transcriptomic structure not only to identify cellular states, but also to reconstruct biologically coherent cell profiles from language-conditioned token generation.

Together, these results show that CellTok preserves the information required to interpret and generate individual cellular states entirely within a token-based autoregressive interface. This bidirectional capacity establishes cellular tokens as native input and output elements of the LLM and provides the foundation for composing multiple cells into structured contexts for subsequent population-level modeling.

### CellTok enables LLMs to interpret cell populations through multi-cell contexts and task semantics

Many downstream analyses in single-cell studies are inherently defined at the level of cell populations, such as cluster annotation, comparative analysis across conditions, and identification of disease-associated states. However, many existing approaches such as LCMs remain fundamentally cell-centric: cells are modeled independently, and population-level conclusions are derived through post hoc aggregation of individual predictions. This paradigm limits the ability to incorporate contextual information arising from population composition, co-occurring transcriptional patterns, and shared biological states across cells.

CellTok addresses this limitation by representing each cell as a compact token sequence and composing multiple tokenized cells within a shared autoregressive context. In this formulation, predictions can be made directly from a multi-cell input, without first reducing the population to independently generated cell-wise outputs. We therefore asked whether CellTok could use both multi-cell transcriptomic context and the semantic information provided by task descriptions to support population-level inference across different settings.

We first evaluated CellTok in a setting that mimics a standard single-cell annotation workflow, in which cells are first grouped into clusters and then assigned labels (**Fig. 2A**). To simulate this process, we constructed homogeneous cell populations from a held-out lung dataset^34^, with each population containing cells from a single cell type and population sizes ranging from 10 to 100 cells (**Methods**). Using donor ctr53 for training and ctr55 for testing, we fine-tuned CellTok-base to assign a shared label to each population, resulting in CellTok-cluster (**Methods**). Because existing methods are not designed to directly annotate cell populations, we adapted them using two strategies: independent per-cell prediction with averaged accuracy, and majority voting across cells in the same population. Although majority voting improved baseline performance, it remained an inherently weaker formulation, as correct prediction only requires the majority of cells to be labeled correctly. As shown in **Fig. 2B**, CellTok outperformed most baselines across different population sizes, while remaining slightly behind scFoundation with majority voting. This result suggests that directly modeling clustered cell populations provides an accurate and scalable solution for realistic annotation workflows.

**Fig. 2.**
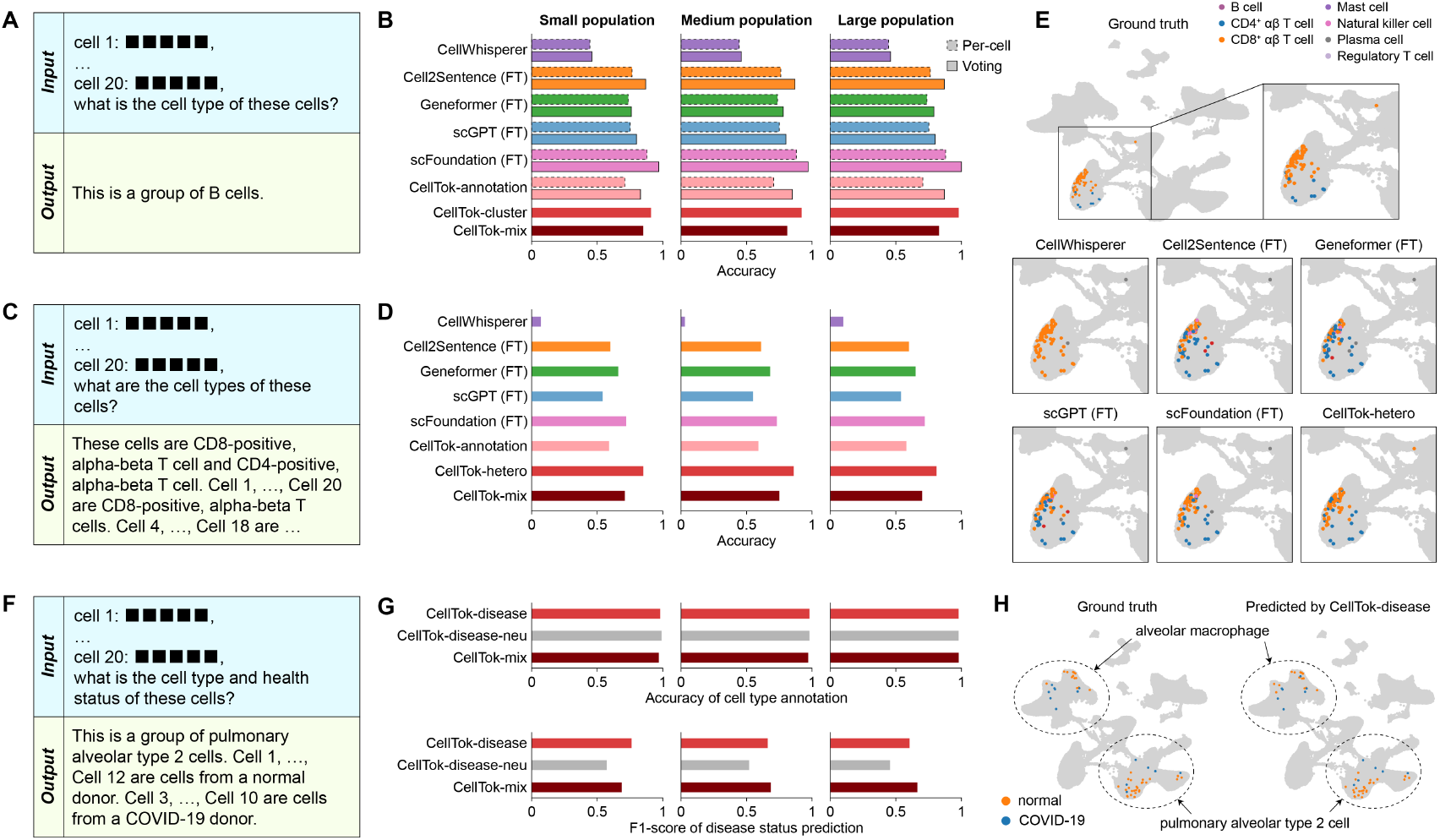
CellTok enables context-aware population-level inference for cell type annotation and disease state identification. (A) Schematic of the cluster-level annotation task, simulating a standard workflow where homogeneous cell populations are first grouped and then assigned a shared label. (B) Performance of CellTok-cluster and CellTok-mix against baseline methods across varying population sizes on the cluster-level annotation task. (C) Schematic of the direct cell annotation task on heterogeneous cell populations, which contain biologically similar and confusable cell types (e.g., CD4^+^ and CD8^+^ αβ T cells) requiring fine-grained resolution. (D) Performance of CellTok-hetero and CellTok-mix compared to baseline methods for direct annotation in heterogeneous populations. (E) Representative UMAP visualization of CD4^+^ and CD8^+^ αβ T cells from the test data, colored by ground truth and predicted cell types. As CellWhisperer cannot be fine-tuned, we aligned its predicted label to the dataset (CD8 T cell to CD8^+^ αβ T cell, and plasma B cell to plasma cell). (F) Schematic of the healthy versus disease-associated state inference task. Cell-type labels are not provided. For CellTok-disease-neu, condition labels (normal and COVID-19) are replaced with semantically neutral labels (Status A and Status B). (G) Performance of CellTok-disease and CellTok-mix in distinguishing healthy from COVID-19-associated cellular states in mixed populations. (H) Representative UMAP visualizations of alveolar macrophages and pulmonary alveolar type II cells from the test data, colored by ground-truth labels and CellTok-disease predictions.

We next asked whether CellTok remains effective when annotation is performed directly on heterogeneous cell populations without relying on purified clusters. To create a more challenging setting, we constructed mixed populations containing two biologically similar and easily confusable cell types, such as CD4^+^ and CD8^+^ αβ T cells, requiring the model to resolve fine-grained distinctions within a shared context (**Fig. 2C**). We fine-tuned CellTok-base on this task to obtain CellTok-hetero (**Methods**). All baseline methods were also fine-tuned, except CellWhisperer, which does not support fine-tuning and was therefore evaluated in a zero-shot setting with label alignment through the unified Hierarchical Annotation Framework (uHAF)^35^ (**Methods**). As shown in **Fig. 2D**, CellTok substantially outperformed all baselines in this direct annotation setting, and this strong performance was maintained even for closely related CD4^+^ and CD8^+^ T-cell populations that are difficult to distinguish (**Fig. 2E**). These results indicate that the contextual modeling framework remains effective even when cell identities must be resolved directly rather than inferred from purified clusters.

Beyond cell-type annotation, many biomedical applications require identifying disease-associated cellular states that are subtle, distributed across multiple cell types, and not reducible to a single marker gene or lineage. This setting presents a different challenge from cell-type annotation: the model must distinguish condition-dependent transcriptional changes within populations, rather than assigning stable lineage identities. To test whether CellTok can capture such state-level variation directly from cellular tokens and population context, we evaluated healthy versus disease-associated state inference without providing cell-type labels at inference time (**Fig. 2F**). Using cells from healthy donor ctr53 and COVID-19 donor cov06 in the lung dataset for training, and ctr55 and cov22 for testing, we constructed populations across nine selected cell types and trained CellTok to identify the disease status of each cell within the population, yielding CellTok-disease (**Methods**). In this setting, cell type labels were not provided during inference, forcing the model to rely on transcriptomic patterns together with population context. CellTok accurately identified disease-associated states across populations constructed from different cell types (**Fig. 2G**). For example, it clearly distinguished disease states in alveolar macrophage and pulmonary alveolar type II cell populations, two lineages closely associated with COVID-19 pathology (**Fig. 2H**). These results indicate that CellTok can reason over population context not only to assign cell identities, but also to detect coordinated disease-related transcriptional programs.

To further examine whether CellTok benefits from the semantic information provided by biological task descriptions, we performed a label-ablation experiment in the disease-state inference task. We replaced the informative condition labels, such as normal and COVID-19, with semantically neutral labels, Status A and Status B, while keeping the underlying transcriptomic inputs and training protocol unchanged (CellTok-disease-neu). This replacement reduced CellTok performance (**Fig. 2G**), indicating that the model does not rely solely on transcriptomic token patterns or supervised label separation. Instead, biologically meaningful labels and task descriptions provide useful semantic priors that help CellTok interpret the inference objective. This result supports the idea that integrating cellular tokens into an LLM interface allows CellTok to leverage language-level biological knowledge when analyzing single-cell populations.

Finally, we asked whether these capabilities could be unified within a single model. We jointly trained CellTok on the three tasks above, including cluster-level annotation, direct cell annotation, and disease-state identification, resulting in a multitask model termed CellTok-mix. Joint training introduced a modest performance trade-off on some task-specific benchmarks, reflecting the increased difficulty of simultaneously optimizing across heterogeneous objectives (**Fig. 2B, 2D, and 2G**). However, CellTok-mix still outperformed most baseline methods on the corresponding evaluation tasks. Notably, its performance even improved on the disease-state identification task, suggesting that shared training across population-level objectives can provide beneficial contextual transfer for relational inference. These results show that the same cellular vocabulary and autoregressive modeling interface can be reused across distinct forms of population-level inference, rather than requiring a separately designed representation or architecture for each task.

### CellTok supports LLM-based inference of cell–cell communication from paired cellular contexts

Beyond assigning identities or states to cells, a central goal of biological analysis is to understand how cells influence each other through intercellular interactions. In single-cell transcriptomics, such interactions are typically manifested as cell–cell communication (CCC), mediated by coordinated ligand–receptor signaling pathways^21^. Unlike cell identity or state, communication is not an intrinsic property of an individual cell, but a relational property conditioned on both interacting cellular contexts. CCC therefore provides a natural setting for testing whether CellTok can predict relational outputs directly from paired cellular inputs, and whether task semantics can help the model map cellular contexts to biologically meaningful communication programs.

We investigated this capability in two complementary settings. First, in a non-spatial setting, we formulated CCC inference as predicting ligand–receptor signaling pathways between a source cell population and a target cell population from their transcriptomic profiles. Using the same held-out lung dataset^34^, we derived putative signaling pathway annotations across healthy and COVID-19 donors with CellChat^36^, and fine-tuned the CellTok-base model, resulting in CellTok-CCC (**Fig. 3A**, **Methods**).

**Fig. 3.**
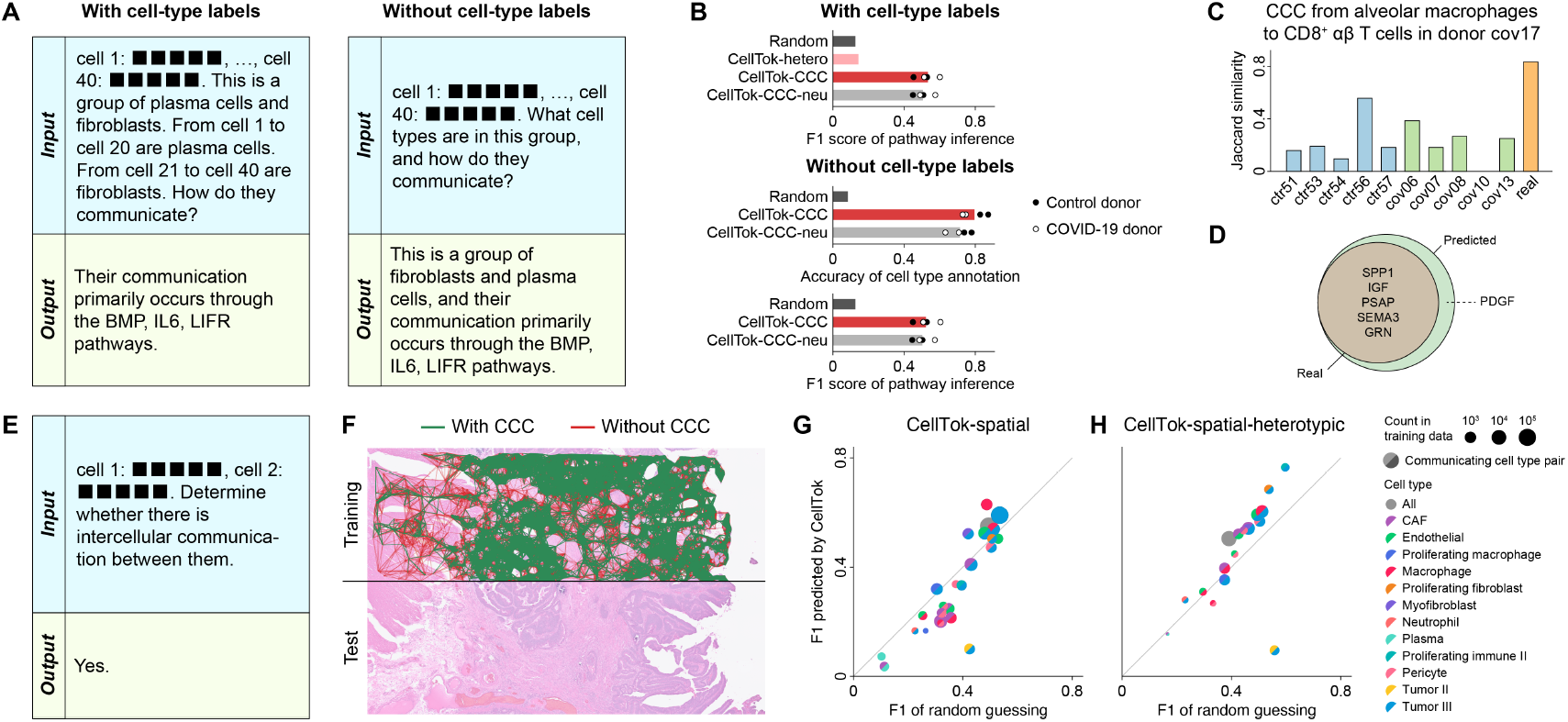
CellTok infers intercellular communication across non-spatial and spatial transcriptomic contexts. (A) Schematic of the non-spatial cell–cell communication (CCC) inference task, where CellTok-CCC predicts active ligand–receptor signaling pathways for source–target cell-type pairs based on transcriptomic profiles with or without predefined cell-type labels. For CellTok-CCC-neu, pathway labels (e.g., BMP, IL6, and LIFR) are replaced with semantically neutral labels (e.g., pathway 1, 2, and 3). (B) Performance of CellTok-CCC in predicting signaling pathways, compared to random guessing and a multiple-choice baseline using CellTok-hetero with only cell-type labels. Bar plots show mean values. (C) Similarity of CellTok-CCC-predicted pathways for donor cov17 to ground-truth pathways from training and test donors, using CCC from alveolar macrophages to CD8^+^ αβ T cells in donor cov17 as an example. (D) Overlap between CellTok-CCC-predicted pathways and ground-truth pathways in donor cov17 for CCC from alveolar macrophages to CD8^+^ αβ T cells. (E) Schematic of the spatial CCC inference task where CellTok-spatial distinguishes communicating from non-communicating spatial neighbors using only transcriptomic input. (F) Human colorectal cancer Visium HD dataset used for spatial CCC inference, showing the selected training region and training CCC pairs. (G–H) F1 scores for individual cell-type pairs predicted by CellTok-spatial (G) or CellTok-spatial-heterotypic (H), compared with a random-guessing baseline that predicts test-set communication events according to the training-set communication ratio of each cell-type pair. Only cell-type pairs with more than 100 training examples in both settings are shown. Circle size indicates the number of training examples. For each circle, the two semicircle colors indicate the two cell types involved in the communication event.

CellTok-CCC achieved strong performance with and without providing cell-type labels (**Fig. 3B**), indicating that CCC signaling pathway inference does not depend on predefined cell identities. In contrast, providing cell-type labels alone is insufficient: CellTok-hetero with cell-type labels under a multiple-choice setting performed only marginally above random guessing (**Fig. 3B, Methods**), suggesting that cell-type information cannot by itself determine communication patterns. Furthermore, pathways predicted by CellTok-CCC were most similar to ground truth from the corresponding test donor rather than to training donors (**Fig. 3C–D, Methods**). This result argues against simple reproduction of training-donor pathway profiles and indicates that the model uses donor-specific transcriptomic information from the paired populations.

We further asked whether the semantic content of pathway names contributes to CCC prediction. To this end, we constructed a semantic-neutralized variant, CellTok-CCC-neu, in which biological pathway names were replaced with uninformative identifiers, such as “pathway 1” and “pathway 2”, while the input cellular contexts and output label structure were kept unchanged. CellTok-CCC-neu showed reduced performance in both cell type annotation and CCC pathway inference compared with CellTok-CCC (**Fig. 3B**). These results suggest that CellTok benefits not only from paired transcriptomic contexts, but also from the biological semantics encoded in task labels. Together, these analyses show that CellTok can predict putative signaling relationships by integrating paired population contexts with biological semantics, rather than relying on cell-type-specific shortcuts or memorized pathway associations.

We next extended this paired-context formulation to spatial transcriptomics, where candidate communication events are constrained by tissue organization. Here, the task was to distinguish communicating and non-communicating cell pairs among spatial neighbors. Notably, CellTok does not take spatial coordinates as input, and must infer communication relationships solely from transcriptomic states (**Fig. 3E**). We evaluated CellTok on a Visium HD human colorectal cancer dataset^37^, where single cells are reconstructed through nuclei segmentation and barcode mapping. Communication labels were derived following the CellNEST^37^ workflow (**Fig. 3F, Methods**).

We selected one tissue region for training and a non-overlapping region for testing. CellTok was trained to classify communicating versus non-communicating neighboring cell pairs using only transcriptomic input, yielding CellTok-spatial (**Methods**). On the held-out region, CellTok-spatial achieved an F1 score of 0.550, outperforming a baseline that predicted test-set communication events according to the training-set communication ratio of each cell-type pair (F1 = 0.495; **Fig. 3G**). This result indicates that transcriptomic states alone contain signals that partially reflect spatial communication relationships. However, stratified analysis by cell-type pair revealed that this improvement was unevenly distributed. CellTok-spatial performed well on cell-type pairs that were highly represented in the training data, most notably Tumor III–Tumor III pairs, which accounted for more than 80% of training examples and showed a substantial improvement over the baseline. In contrast, performance on less frequent cell-type pairs was weaker and, in some cases, fell below the baseline, suggesting that the model was partly influenced by the dominant homotypic interaction pattern.

To test whether CellTok could learn more general communication signatures across different cell types, we further removed homotypic interactions and constructed a dataset consisting only of heterotypic neighboring cell pairs. We trained a corresponding model, CellTok-spatial-heterotypic, under the same spatial generalization setting (**Methods**). Although its overall F1 improvement over the cell-type-pair baseline was comparable to that of CellTok-spatial, CellTok-spatial-heterotypic showed stronger-than-baseline performance across most cell-type pairs rather than being driven by a single dominant pair (**Fig. 3H**). Importantly, cell-type labels were not provided to the model during CCC prediction. Thus, the improved performance across diverse heterotypic pairs suggests that CellTok does not simply memorize predefined cell-type identities or pair frequencies, but instead captures transcriptomic features associated with communication competence across cellular contexts.

Together, these results show that CellTok can use paired cellular contexts to predict two forms of CCC-related output: signaling pathways between non-spatial cell populations and communication status between spatially neighboring cells. Although both tasks use supervision derived from computational CCC methods, they provide proof-of-concept evidence that the same token-based interface can represent and predict relational properties between cells and cellular populations. The pathway-name ablation further suggests that embedding cellular tokens within a language-model interface enables CellTok to combine molecular context with task semantics, providing a potential advantage over models that treat biological labels as arbitrary categorical outputs.

### CellTok enables LLMs to resolve developmental ordering from cellular tokens

A central property of cellular systems is their continuous evolution over time. We thus asked whether CellTok can recover temporal structure directly from transcriptomic profiles by inferring the developmental ordering of cellular states. We evaluated this capability using a scRNA-seq dataset of brain organoids derived from three induced pluripotent stem cells (iPSCs) and one embryonic stem cell (ESC)^38^, spanning a developmental trajectory from day 4 to 2 months and covering multiple stages of differentiation (**Fig. 4A**).

**Fig. 4.**
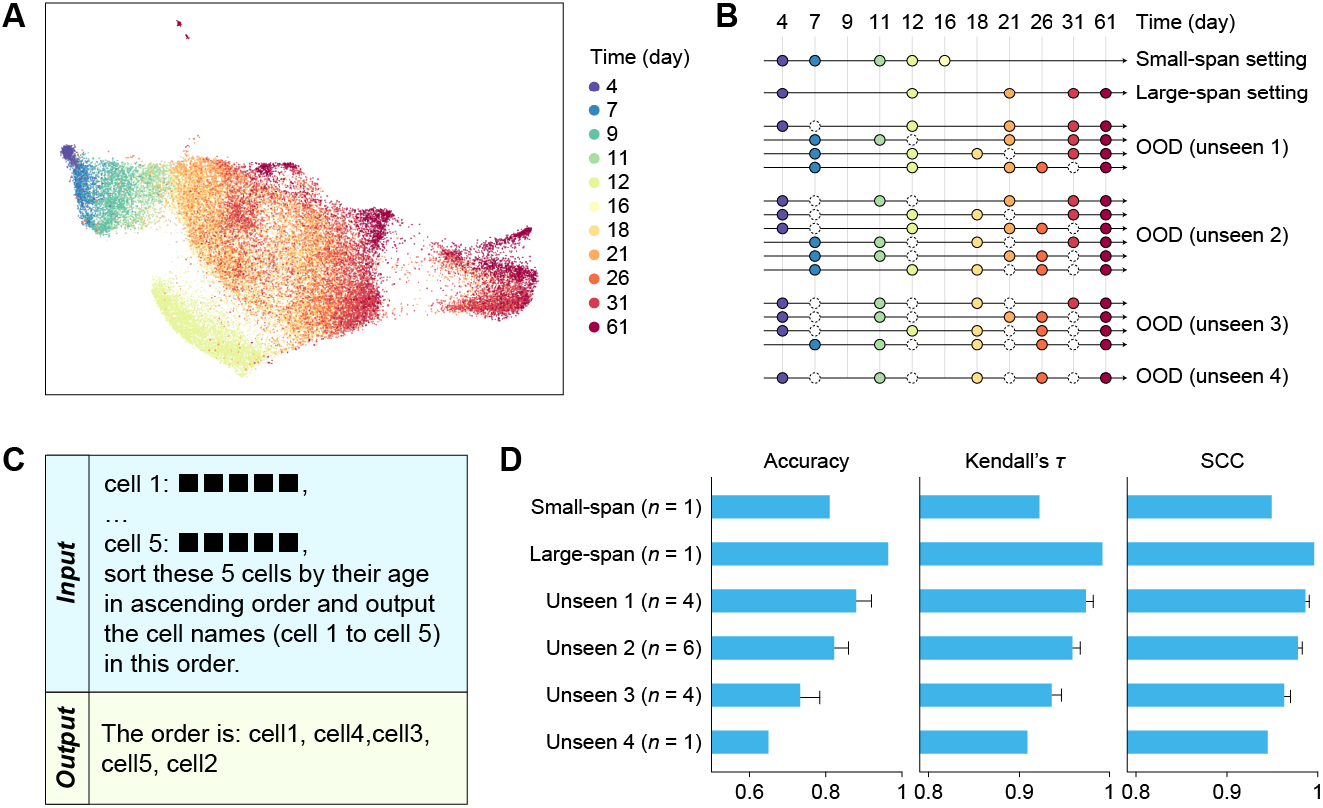
Inferring pseudotime along developmental trajectories with CellTok. (A) UMAP visualization of the brain organoid dataset, illustrating the developmental trajectory of cells derived from induced pluripotent stem cells (iPSCs) and embryonic stem cells (ESCs) across multiple stages of differentiation spanning from day 4 to day 61. (B) Experimental design for trajectory inference, detailing two temporal resolution settings: a small-span setting (densely sampled, transcriptionally similar stages) and a large-span setting (widely separated time points). The schematic also illustrates the out-of-distribution (OOD) experimental setup, where observed time points are replaced with unseen days. (C) Schematic of the pseudotime inference task. (D) Performance of CellTok on pseudotime inference across different temporal resolution settings. Bar plots show mean values, with error bars representing the standard error of the mean (SEM). Kendall’s *τ*: Kendall rank correlation coefficient; SCC: Spearman correlation coefficient.

To examine temporal ordering at different resolutions, we designed two settings (**Fig. 4B**): a large-span setting with widely separated time points (days 4, 12, 21, 31, and 61), and a small-span setting with more densely sampled and transcriptionally similar stages (days 4, 7, 11, 12, and 16). In both settings, CellTok was trained with sampled developmental days as supervision and asked to infer the relative temporal positions of held-out cells from their transcriptomic profiles. Performance was evaluated using accuracy, Kendall rank correlation coefficient, and Spearman correlation coefficient (**Fig. 4C, Methods**).

Across both settings, CellTok achieved high temporal ordering performance (**Fig. 4D**). As expected, performance was stronger in the large-span setting, where developmental stages are more transcriptionally separated. Importantly, CellTok also maintained reliable performance in the small-span setting, indicating that it could resolve relatively fine-grained temporal differences among nearby developmental stages rather than only separating distant cell states.

To assess whether CellTok learned temporal relationships that generalized beyond the exact time points used during training, we performed an out-of-distribution (OOD) local temporal generalization experiment in the large-span setting. We replaced increasing numbers of training-observed days with nearby but unseen days while preserving the overall ordering task (**Fig. 4B**). Performance gradually decreased as more time points were substituted, but remained robust even when four of the five stages were replaced (**Fig. 4D**). This result suggests that CellTok did not simply memorize the sampled day labels, but learned transcriptomic patterns that remain locally consistent along developmental progression.

Together, these results show that CellTok can infer the relative ordering of cellular states across developmental stages and generalize this ordering to nearby unseen time points. This extends the token-based modeling interface beyond static cellular populations to the temporal organization of cellular states.

### CellTok enables LLMs to generate stage-matched transcriptomic profiles from developmental contexts

Having shown that CellTok can recover the relative ordering of cellular states across developmental stages, we next asked whether the same token-based formulation could support conditional generation along a developmental trajectory. Specifically, we evaluated whether CellTok could generate transcriptomic profiles matching a target developmental stage from cells sampled at other stages, and whether this capability generalized across different temporal configurations and generation regimes (**Fig. 5A**).

**Fig. 5.**
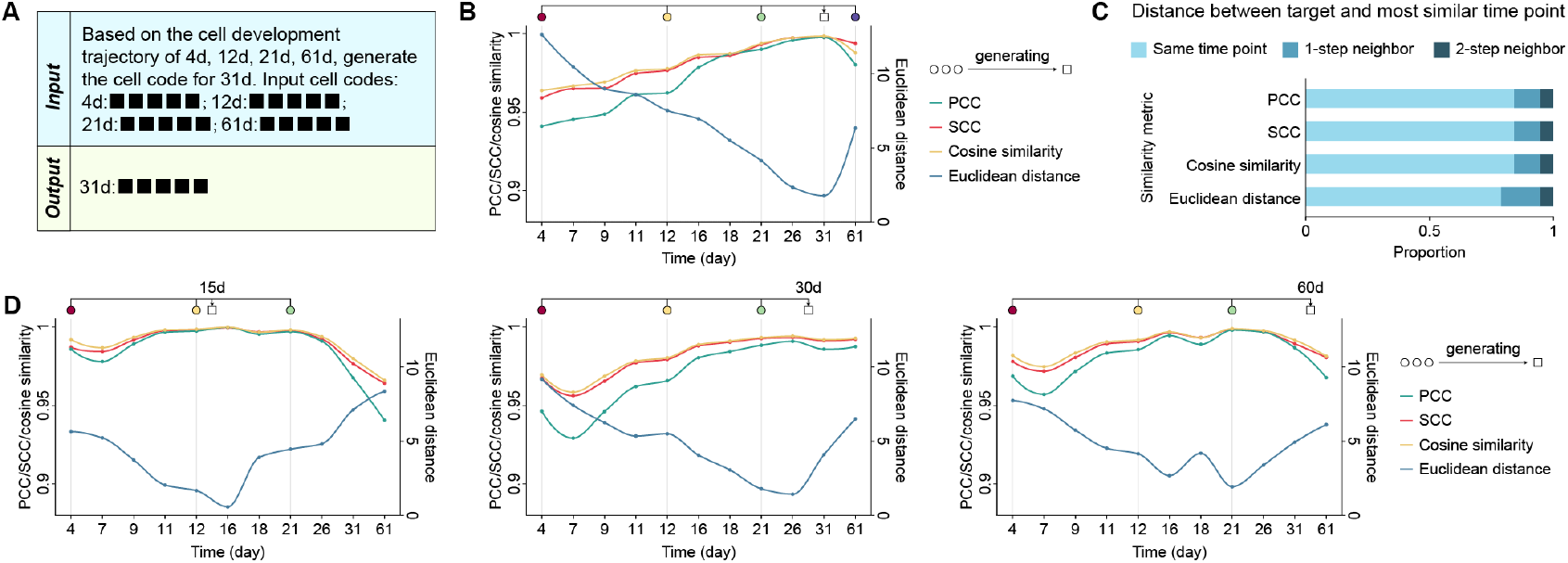
Generating gene expression profiles along developmental trajectories with CellTok. (A) Schematic of the trajectory-conditioned generation task. (B) Performance of the fine-tuned CellTok model in the setting where CellTok is prompted to generate target cells at day 31 using contextual cells sampled from days 4, 12, 21, and 61. (C) Performance of CellTok-trajectory across all generation settings (*n* = 19). For each generated cell, the most similar real time point was identified using different similarity metrics, and its distance from the target time point was then calculated. (D) OOD evaluation for unseen target days. PCC: Pearson correlation coefficient; SCC: Spearman correlation coefficient.

We first considered a base setting in which the model was fine-tuned to generate day 31 cells from cells collected at days 4, 12, 21, and 61 (**Fig. 5B**). In this proof-of-concept setting, generated cells were most similar to real cells from the target day, indicating that CellTok could produce transcriptomic profiles consistent with the intended developmental stage from temporally distributed cellular contexts.

We next constructed a diverse set of generation settings that varied temporal spacing and prediction regime, including local interpolation, long-range interpolation, and extrapolation toward trajectory boundaries (**Methods**). We fine-tuned CellTok across these settings to obtain CellTok-trajectory and evaluated generation fidelity by comparing mean expression profiles between generated and real cells using complementary correlation- and distance-based metrics (**Methods**). Across most settings, generated profiles showed the highest agreement with real profiles from the corresponding target time points (**Fig. 5C**), suggesting that CellTok captured a trajectory-consistent mapping rather than overfitting to a single temporal configuration.

We further evaluated CellTok-trajectory in an unseen target-day setting, in which the requested target days were absent from training (**Methods**). In this setting, the model was required to generate cells at temporally nearby but previously unobserved developmental stages. As the temporal distance between the target day and the conditioning stages increased, generation accuracy gradually declined. Nevertheless, the real time points most similar to the generated profiles remained close to the intended targets (**Fig. 5D**), suggesting that CellTok captured local continuity in developmental-stage-associated transcriptomic variation rather than merely reproducing discrete target days observed during training.

Together, these results show that CellTok supports developmental-stage-conditioned generation across varying temporal contexts and can generalize to nearby unseen target days. This extends the native token-based interface from temporal ordering to conditional generation of cellular states along developmental progression.

## Discussion

In this work, we propose CellTok, a tokenized single-cell language modeling approach that enables LLMs to read transcriptomic profiles as discrete cellular tokens. By converting continuous expression profiles into compact token sequences, CellTok provides a language-compatible cellular representation and an LLM-native modeling interface through which individual cells can be arranged, composed, predicted, and generated within structured cellular contexts. This formulation moves single-cell modeling beyond treating cells primarily as endpoints of molecular representation, and instead makes cellular states native, composable elements in the token language of LLMs for modeling populations, intercellular relationships, and developmental processes.

A key implication of CellTok is that diverse population-level analyses can be reformulated as sequence modeling problems within LLMs. Instead of relying exclusively on downstream aggregation of cell embeddings or separately designed task-specific architectures, CellTok allows multiple cells, biological descriptions, and relational outputs to be represented within a shared token space and processed through the same autoregressive interface. Population annotation, disease-state inference, cell–cell communication prediction, trajectory reasoning, and stage-conditioned generation can therefore be expressed as different conditional prediction tasks over cellular and textual tokens.

The integration of cellular and textual tokens also creates a direct interface between molecular measurements and natural language in scientific research. In CellTok, cellular tokens can condition textual descriptions, while textual descriptions can condition the generation and interpretation of cellular states. This bidirectional grounding makes cellular tokens usable not only as internal representations, but also as input and output elements of language-model-based analysis. Importantly, our experiments suggest that CellTok can benefit from task-relevant biological information expressed in natural language. These results provide initial evidence that cellular tokenization may allow LLMs to use biological task semantics and prior knowledge when analyzing cellular contexts.

This capability is central to the motivation of CellTok. Many biological questions require not only recognizing molecular patterns, but also relating them to phenotypes, pathways, conditions, and mechanistic hypotheses. Conventional LCMs provide powerful representations of cellular states, but these representations often remain external to the linguistic space in which biological knowledge and scientific questions are formulated. By making cellular states available as tokens within the same modeling space as biological language, CellTok provides a route for connecting molecular measurements with knowledge-guided inference. The current study demonstrates this possibility at an early stage, and systematic evaluation of knowledge usage and reasoning behavior will be important for future work.

CellTok should therefore be viewed as a proof of concept for cellular tokenization as a shared representation and task interface, rather than as a task-specific model optimized for a single benchmark or as a single parameter set that solves all single-cell problems. Most analyses in this study use task-specific fine-tuning, while CellTok-mix provides an initial demonstration that the same cellular vocabulary and autoregressive interface can be reused across several related population-level tasks. Notably, joint training introduced only a modest performance trade-off in most settings and even improved performance in the disease-state identification task, suggesting that some forms of population-level reasoning may benefit from shared training across related cellular contexts.

Several limitations and future directions remain. First, although discretization provides compactness and compositionality, it may compress or discard fine-grained quantitative information from continuous gene expression profiles. Future work should characterize which biological signals are preserved or lost during tokenization and develop tokenizers that better balance compression, interpretability, and quantitative fidelity. Second, the semantics and transferability of learned cellular tokens depend on the training data, gene panel, codebook design, and tokenizer architecture. More expressive tokenization schemes, including tokenizers initialized from or built upon pretrained cellular representations from LCMs, may improve token fidelity and transferability while reducing dataset-specific biases. Third, although CellTok supports several tasks through a shared token-based interface and can benefit from task-relevant prompts, its ability to transfer knowledge across tasks and biological contexts remains to be systematically evaluated. Larger multitask corpora, more explicit instruction tuning, and systematic transfer experiments will be needed to determine whether knowledge learned in one cellular context can improve performance in another. Finally, cellular tokenization can be extended beyond transcriptomics to incorporate chromatin accessibility, proteomics, perturbation profiles, and spatial information, enabling richer cellular tokens for multi-omic and tissue-level modeling.

More broadly, CellTok points to a possible route toward LLM-based science in single-cell biology. For LLMs to contribute to discovery, they must be able to operate not only on scientific text, but also on the biological objects that science seeks to understand. By making cellular states readable within the same token space as biological language, CellTok provides an initial attempt to bring single-cell data into the native language of LLM-based scientific reasoning, enabling future models to interpret molecular measurements in closer connection with biological knowledge and scientific questions.

## Methods

### Cell tokenizer

#### Model architecture

Inspired by VQ-VAE^23^, we developed a discrete tokenization model that maps each cell’s gene expression profile to a compact sequence of discrete codes. For an input cell-by-gene expression matrix *x*, all datasets were first aligned to a shared panel of 5,000 genes. Raw expression counts were normalized using library-size normalization followed by log transformation:

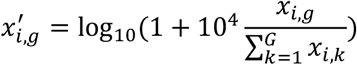

where *x*_*i,g*_ denotes the raw expression of gene *g* in cell *i*, and *G* is the total number of genes.

The tokenizer consists of an encoder, a vector quantization module, and a decoder. The encoder maps each normalized expression profile 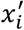 to a continuous latent representation 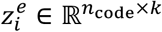, which is then partitioned into *n*_code_ segments 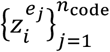,each of dimensionality *k*. The vector quantization module maintains a learnable codebook 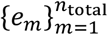, where each codebook vector *e*_3_ ∈ ℝ^k^

Each encoder segment 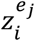 is then replaced by its nearest codebook vector according to Euclidean distance:

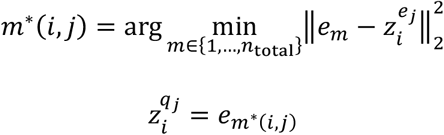

where 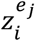 is the *j*-th segmentation of the *i*-th cell’s embedding, and 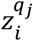 is the segmentation after quantization.

The quantized latent representation 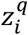 is obtained by concatenating all quantized segments:

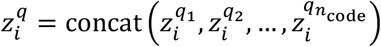

The decoder reconstructs the normalized expression profile 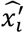 from 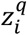. To adapt to the characteristics of scRNA-seq data, we employed LayerNorm and PReLU activations in both the encoder and decoder.

The tokenizer was trained by minimizing the following loss:

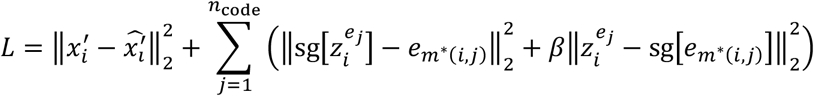

where sg[·] denotes the stop-gradient operator, which prevents gradient flow through the indicated tensor. The first term corresponds to the reconstruction error, for which we used mean squared error (MSE) between the reconstructed and normalized inputs. The remaining terms encourage the encoder outputs to commit to and align with the learned codebook vectors. β is a hyperparameter controlling the strength of the commitment loss.

To train the discrete cell tokenizer, we curated approximately 68 million single-cell expression profiles from CELLxGENE^25^ after quality filtering, spanning 285 tissues and 660 refined cell types across healthy and diseased conditions. These datasets were split at the study level into 63.8 million training cells and 4.7 million test cells. The tokenizer was trained solely on normalized gene expression profiles without using any textual or metadata information.

#### Experiments for determining parameters

To determine suitable hyperparameters for the tokenizer, we performed ablation experiments on three key components: the total number of codebook vectors (*n*_total_), the number of code segments (*n*_code_), and the dimensionality of each code vector (*k*). We first fixed *n*_code_ = 32 and varied *n*_total_ and *k*, and then fixed the total latent dimensionality (*n*_total_ = 256) to examine the effects of different values of *n*_code_ and *k*.

All tokenizer variants were trained on the training split and evaluated on the held-out test set. Performance was assessed using metrics in both the latent space and the reconstruction space, including MSE, Pearson correlation coefficient (PCC), average cosine similarity (ACS), adjusted Rand index (ARI), normalized mutual information (NMI), and adjusted mutual information (AMI).

As shown in **Fig. S1A–C**, reducing the number of code segments (*n*_code_) led to a pronounced degradation in both latent and reconstruction performance, whereas variations in *n*_total_ and *k* had comparatively minor effects. Considering both empirical performance and representational efficiency, we selected *n*_total_ = 256, *n*_code_ = 32, and *k* = 8 as the default tokenizer configuration.

#### Experiments for determining gene panels

We evaluated the reconstruction performance of the tokenizer using different gene panels, including 2,000 highly variable genes (HVGs), 5,000 HVGs, and an 18,980-gene panel covering most protein-coding genes. As shown in **Fig. S1D**, the 5,000-gene panel consistently achieved the best reconstruction performance across evaluation metrics. Based on this empirical comparison, we selected the 5,000-gene panel as the default setting for all experiments.

### Integrating cell tokens into the LLM

#### Training strategy

After training the tokenizer, we incorporated the *n*_total_ learned codebook entries as cell tokens into the vocabulary of a pretrained LLM (Qwen 2.5 7B^24^). Each codebook entry was assigned a unique token ID, treated identically to standard text tokens, enabling joint autoregressive training.

Given a mixed token sequence *x* = (*x*_1_,…,*x*_*T*_) composed of text tokens and cell tokens, the model was trained to maximize the autoregressive log-likelihood:

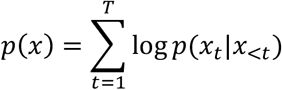

where *x*_<*t*_ denotes the preceding tokens.

To efficiently adapt the pretrained LLM, we employed Low-Rank Adaptation (LoRA)^39^, which introduces trainable low-rank matrices into the frozen linear layers and vocabulary projection layers. Each adapted linear transformation is defined as:

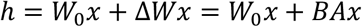

where *W*_0_ is the frozen parameters of Qwen, and *A* ∈ ℝ^D×E^,*B* ∈ ℝ^E×D^ are trainable matrices with rank *r* ≪ *d*, providing a parameter-efficient mechanism for model adaptation.

For all CellTok tasks, we fine-tuned the pretrained Qwen model using LoRA by adapting the attention projection layers, feed-forward layers, token embedding layers, and the output token prediction projection layers. This resulted in training approximately 2_1_ of the total parameters of the Qwen backbone.

#### Ablation across LLM backbones

To evaluate the robustness of CellTok across different LLM backbones, we performed experiments using three LLMs of varying scales and architectures: Qwen 2.5 0.5B, Qwen 2.5 3B, and LLaMA 3 8B. For each backbone, the vocabulary was extended with the same set of learned cell tokens, and models were fine-tuned using the same training protocol.

Results are shown in **Fig. S1E**. Performance consistently improved with increasing model scale, in line with expected LLM scaling behavior. Importantly, comparable trends were observed across different model families, indicating that the proposed tokenization and early-fusion strategy generalize beyond a specific LLM backbone.

#### Tasks for training CellTok-base

To fine-tune the LLM to obtain CellTok-base, each cell was first tokenized using the tokenizer and represented as a short sequence of discrete cell tokens. To enable the pretrained LLM to understand, generate, and reason over these cell tokens, we formulated two complementary pretraining tasks for CellTok that jointly couple cellular and linguistic representations.

In the comprehension task, the model is conditioned on a sequence of cell tokens corresponding to a single cell and is trained to generate a natural-language description of the cellular state. This task encourages the model to map discrete cell tokens to semantically meaningful textual concepts such as cell identity. In the generation task, the model is conditioned on a textual description of a cell and is trained to generate the corresponding sequence of cell tokens. This task enables the model to synthesize discrete cellular representations from language, which can subsequently be decoded into gene expression profiles by the tokenizer decoder.

Together, these two tasks align cellular and linguistic modalities within a unified autoregressive framework, encouraging bidirectional grounding between cell tokens and natural language. During pretraining, both tasks were optimized jointly using standard next-token prediction, with task-specific prompts determining whether the model was trained in the comprehension or generation mode.

For training the base model of CellTok, we used the same single-cell expression profiles as those used for tokenizer training. Cell metadata were converted into descriptive text using GPT-4o^26^, yielding paired cell token–text data for multimodal training.

To preserve the general language understanding ability of the pretrained LLM during multimodal adaptation, we further mixed these cell token–text samples with general text samples at a 1:1 ratio during training. This mixed training strategy allowed the model to acquire cellular token understanding while maintaining its original linguistic competence.

### Cell-level annotation and generation tasks

We evaluated CellTok against representative baseline methods on cell-level annotation and generation tasks. For methods that support fine-tuning (scGPT, Geneformer, scFoundation, Cell2Sentence, and CellTok), each dataset was split into 10,000 training cells and 1,000 testing cells. CellTok was fine-tuned separately on each dataset to obtain the task-specific model CellTok-annotation. All baseline models under the fine-tuning setting were trained for five epochs using the default hyperparameters recommended in their official tutorials.

For general-purpose LLMs, we followed the evaluation protocol of GPTCelltype^17^, adapting the corresponding APIs to each model. To account for differences in label granularity across datasets and methods, predicted labels were aligned using the unified Hierarchical Cell Annotation Framework (uHAF)^35^. CellWhisperer, which does not provide fine-tuning code, was evaluated only under the zero-shot setting. Performance on the annotation task was assessed using accuracy and macro-F1 score.

For the gene expression generation task, all models were trained and evaluated using the same data splits as in the annotation task, and were configured using default parameters from their official implementations.

### Population-level annotation tasks

We evaluated population-level cell annotation using a held-out lung dataset^34^. For the cluster-level annotation task, to mimic a standard single-cell annotation workflow, in which cells are first grouped into clusters and then annotated at the cluster level, we performed Leiden clustering on the dataset. For each Leiden cluster, we calculated the proportion of cells belonging to each annotated cell type. Clusters in which a single cell type accounted for more than 90_1_ of cells were considered homogeneous clusters. Cell populations were then constructed by randomly sampling cells from these homogeneous clusters, with the dominant cell type of the source cluster used as the population-level label. One healthy donor (ctr53) was used for training, and a different healthy donor (ctr55) was used for testing to ensure donor-level separation. To assess scalability with respect to population size, we defined three population-size regimes: small (10–20 cells), medium (20–50 cells), and large (50–100 cells). We randomly sampled 2,000 cell populations from the training donor for model fine-tuning. For evaluation, we sampled 100 cell populations from the held-out test donor for each population-size regime.

For all methods that support fine-tuning, models were trained on the training populations for five epochs using the default hyperparameters specified in their official implementations. For CellTok-cluster, predictions were made directly at the population level, and accuracy was computed based on the predicted shared cell type for each group. For baseline methods that operate only at single-cell resolution, we adopted two evaluation strategies. In the without-voting strategy, cell types were predicted independently for each cell within a population, and group-level accuracy was obtained by averaging cell-level accuracies within each group. In the majority voting strategy, the most frequently predicted cell type within a population was taken as the group-level prediction, and accuracy was computed accordingly. For zero-shot evaluations, predicted labels were mapped using uHAF prior to metric computation, whereas fine-tuned models were evaluated directly without additional label alignment.

We further evaluated population-level annotation under a more challenging heterogeneous setting, in which each population contains a mixture of two different cell types. Using the same training and testing donors, we selected seven cell types and intentionally included biologically similar and commonly confounded pairs, such as CD4^+^ and CD8^+^ αβ T cells. From the training set, we randomly sampled 2,000 mixed populations, and from the test set, 100 mixed populations. Within each population, cells from the two types were randomly interleaved to avoid ordering biases. The same three population-size regimes were used as in the homogeneous setting.

For evaluation, predictions were assessed without voting by computing cell-level predictions within each population and aggregating performance across groups. Metrics were computed by comparing the predicted and ground-truth cell-type assignments within each population, and final results were obtained by averaging across all test populations.

### Disease state identification task

We evaluated disease state identification at the population level using the same lung dataset^34^. One healthy donor (ctr53) and one COVID-19 donor (cov06) were used for training, while a different healthy donor (ctr55) and COVID-19 donor (cov22) were reserved for testing to ensure donor-level separation. We focused on nine selected cell types and defined three population-size regimes based on the total number of cells in each group, consistent with the population-level annotation tasks.

Cell populations were constructed by randomly sampling cells from both healthy and COVID-19 donors such that each population contained a mixture of disease states. Within each population, cells were drawn from a single cell type, and the ordering of healthy and diseased cells was randomly shuffled to avoid positional bias. We generated 2,000 training populations and 100 test populations for each population-size regime.

The model was evaluated under the fine-tuned setting. Disease state prediction was performed at the cell level within each population, and performance was summarized using group-level accuracy aggregated across test populations.

### Multi-task training of CellTok-mix

To train CellTok-mix, we combined datasets from homogeneous population annotation, heterogeneous population annotation, and disease state inference into a unified corpus. The CellTok-base model was then fine-tuned on this aggregated dataset to obtain CellTok-mix. During this fine-tuning, we also mixed the task-specific training samples with general text samples to preserve the language understanding ability of the pretrained LLM.

### Non-spatial CCC inference task

For the non-spatial CCC inference task, we used the lung dataset^34^. Five healthy donors (ctr51, ctr53, ctr54, ctr56, ctr57) and five COVID-19 donors (cov06, cov07, cov08, cov10, cov13) were used for training, while two healthy donors (ctr55, ctr52) and two COVID-19 donors (cov22, cov17) were reserved for testing to ensure donor-level separation.

We focused on 12 commonly shared cell types across donors. To obtain supervision signals for CCC, we applied CellChat^36^ independently to each donor to infer potential ligand– receptor–mediated signaling pathways between all pairs of cell types. For each source–target cell type pair, we randomly constructed 20 cell population pairs, where each population pair consisted of 20 source cells and 20 target cells sampled from the corresponding donor. These population pairs were formatted into prompts and used to train CellTok-CCC. This formulation treated the inference as a generative completion task, where the model was required to complete the sequence of communication pathways bridging two given cell populations. Model predictions were evaluated using pathway-level precision, recall, and F1 score.

For the zero-shot evaluation of CellTok-hetero, we provided a candidate pool consisting of all 73 unique pathways appearing in the dataset. The model was tasked with identifying the correct subset of pathways from this comprehensive list, effectively framing the inference as a multiple-choice selection task.

As for the baseline, we employed a random sampling strategy that selected a subset from the 73-pathway pool, matching the number of pathways predicted by CellTok-hetero, to evaluate performance against the ground truth.

To assess whether the model infers communication pathways based on population-specific transcriptomic features rather than memorizing source–target cell type identities, we performed an additional analysis on two held-out cases from donor cov17. We computed the Jaccard similarity between the model-predicted pathways and (i) the ground-truth pathways of the test donor and (ii) pathway sets observed in the training donors. This analysis evaluated whether predictions are more consistent with the test donor–specific communication context than with training donor patterns.

### Spatial CCC inference task

For the spatial CCC inference task, we processed the human colorectal cancer Visium HD dataset^37^ following the CellNEST workflow^37^. Single cells were reconstructed by integrating nuclei segmentation with Visium HD barcode information, and candidate CCC events were defined among spatially neighboring cells based on ligand–receptor co-expression.

Specifically, for a potential communication event from cell *i* to cell *j*, cell *j* was required to be among the top 50 spatial neighbors of cell *i* based on Euclidean distance. A ligand–receptor interaction was considered active when the ligand in the source cell and the receptor in the target cell both exceeded the 98.5th percentile of their respective cell-specific gene expression profiles, corresponding to the top 1.5_1_ expressed genes in each cell. Cell pairs satisfying both the spatial proximity and ligand–receptor co-expression criteria were labeled as positive communication pairs. To construct a balanced classification dataset, we generated an equal number of negative samples from spatially neighboring cell pairs that did not satisfy the ligand–receptor co-expression criterion. This design ensured that both positive and negative samples were drawn from spatially proximal cell pairs, so that the task focused on distinguishing communication-related transcriptomic patterns rather than simply separating nearby from distant cells.

To evaluate spatial generalization, we split the tissue section into two non-overlapping regions. Cell pairs located in one region were used for model training and validation, whereas cell pairs from the other region were held out for testing. This region-level split was designed to assess whether the model could generalize communication prediction to an unseen spatial area rather than relying on local spatial neighborhoods. CellTok-spatial was trained to predict whether each neighboring cell pair represented a communication event using only transcriptomic input, without using spatial coordinates or cell-type labels.

To further evaluate whether CellTok could learn communication patterns beyond dominant homotypic interactions, we constructed an additional heterotypic CCC dataset by removing cell pairs in which the source and target cells belonged to the same annotated cell type. This dataset retained only neighboring cell pairs between distinct cell types and was split using the same spatial generalization strategy. The heterotypic dataset was used to train CellTok-spatial-heterotypic. This setting specifically tested the model’s ability to infer communication events across different cell-type contexts from transcriptomic states alone.

To evaluate whether CellTok learned communication-related transcriptomic signals beyond cell-type-pair frequency, we constructed a random-guessing baseline based on the training-set communication ratio of each cell-type pair. For each ordered cell-type pair, we first calculated the positive ratio in the training set, defined as the fraction of neighboring cell pairs labeled as communicating among all training pairs of that cell-type combination. For each test cell pair belonging to the same cell-type pair, the baseline then randomly predicted it as positive with this training-set positive ratio, and as negative otherwise. Thus, this baseline uses cell-type-pair-level communication frequency from the training data, but does not use transcriptomic profiles of individual cells. Because the baseline prediction is independent of the true label for each test pair, its expected recall equals the positive ratio in the training set, whereas its expected precision equals the positive ratio in the test set. The corresponding F1 score was calculated from these expected precision and recall values. This baseline therefore represents the performance expected from knowing only how frequently each cell-type pair communicates in the training region, without learning cell-state-specific communication patterns.

### Pseudotime inference tasks

Using the brain organoid dataset^38^, we defined two temporal configurations to probe trajectory inference at different resolutions: a small-span setting (Days 4, 7, 11, 12, 16) and a large-span setting (Days 4, 12, 21, 31, 61). For each setting, we randomly sampled 20,000 distinct cell sequences for training and 5,000 for evaluation.

To probe generalization beyond observed time points, we constructed OOD test sets by systematically replacing each time point with a nearby, unobserved day. Specifically, Day 4 was replaced with Day 7, Day 12 with Day 11, Day 21 with Day 18, and Day 31 with Day 26. This setup tested whether the model captures continuous temporal structure rather than memorizing discrete sampling points.

### Gene expression generation along developmental trajectories

Using the brain organoid dataset^38^, we formulated trajectory-conditioned generation as a temporal imputation problem along developmental trajectories. Given cell code sequences from a set of observed time points (context), the model was trained to generate the gene expression profile at a held-out target day, enabling both interpolation and extrapolation along the timeline.

We constructed a mixed suite of 19 generation tasks (total *n* = 260,000), grouped into four regimes, to train CellTok-trajectory. Each task was defined as predicting a target day from a set of context days: three local interpolation tasks used nearby temporal neighbors as context, where the target lies within the local neighborhood (e.g., context Days 4, 9, and 11 with target Day 7); two dense interpolation tasks incorporated richer multi-point context (e.g., context Days 4, 7, 9, 12, and 16 with target Day 11); five mid-range interpolation tasks placed the target between temporally distant anchors (e.g., context Days 4, 12, and 21 with target Day 16); nine extrapolation tasks predicted early or late endpoints from boundary observations (e.g., context Days 4, 7, and 9 with target Day 11). For each task, we performed a stratified split with a 7:2:1 ratio for training, validation, and testing, and then merged splits across tasks, yielding 182,000 training, 52,000 validation, and 26,000 test instances in total.

To assess whether CellTok-trajectory can generalize beyond task definitions observed during training, we evaluated the trained model under a target-day out-of-distribution (OOD) setting. In this setting, the conditioning set was kept fixed, while the target day was replaced with an unseen temporal node. For example, using context cells from Days 4, 12, and 21, the model was required to generate cells for Day 15 rather than the training-observed target Day 16. For each OOD task, we randomly sampled 5,000 instances to construct a dedicated test set. The model was evaluated directly on these held-out tasks without further fine-tuning.

## Data availability

Data used for pre-training CellTok were obtained from CELLxGENE^25^, which can be downloaded from https://cellxgene.cziscience.com/. All held-out single-cell datasets for validation are contained in the CELLxGENE dataset. The Visium HD human colorectal cancer dataset can be downloaded from https://www.10xgenomics.com/datasets/visium-hd-cytassist-gene-expression-libraries-of-human-crc. The brain organoids dataset can be downloaded from https://www.ebi.ac.uk/biostudies/arrayexpress/studies/E-MTAB-12001.

## Code availability

The source code for tests and evaluations is available online on GitHub at https://github.com/sunnsset/CellTok.

## Acknowledgments

The work is supported in part by the National Natural Science Foundation of China (grants 92470105, 62373210), the National Key R&D Program of China (grants 2025YFC3409300, 2024YFF0729200), and Tsinghua-Toyota Joint Research Institute Inter-disciplinary Program (20243930093).

## Author contributions

X.Z., L.W., C.X., and H.B. conceived the study. X.Z. and L.W. supervised the study. C.X., Y.D., and H.B. designed the model, collected the data, designed the experiments and performed the analysis. Y.C. contributed to data analysis. X.Z., L.W., C.X., and Y.D. wrote the manuscript. All authors read and approved the final manuscript.

## Declaration of interests

The authors declare no competing interests.

## Notes

### Competing Interest Statement

The authors have declared no competing interest.

### Summary of Updates

This revised version of the manuscript includes the following major updates compared with the previous version: 1. Title and Abstract: The title has been rephrased to more accurately reflect the scope and main findings of the study. The abstract has been rewritten to better summarize the research background, methods, key results, and conclusions. 2. Overall framing and conception: The overall framing and narrative of the manuscript have been substantially revised. The Introduction has been reorganized to clarify the scientific question, highlight the knowledge gap, and better position the study within the current literature. The logical flow between sections has also been improved to strengthen the central message of the paper. 3. Additional experiments: New experiments have been added to further support the main conclusions and address potential concerns regarding the robustness of the results. The corresponding data, figures, and descriptions have been incorporated into the Results section. 4. New sections and expanded content: Several sections have been added or expanded to provide a more complete presentation of the work, including additional methodological details, extended analyses, and further discussion of the biological implications of our findings. 5. Figures and tables: New figures and supplementary items have been added to present the newly included experimental results. Existing figures have been updated where necessary to maintain consistency with the revised text. 6. Discussion: The Discussion section has been revised to incorporate the new results, discuss their implications in a broader context, and address the limitations of the study. 7. References and language: The reference list has been updated to include recently published relevant work. The language of the manuscript has also been polished throughout to improve clarity and readability.

